# Ophiobolin A selectively alters mitochondria, metabolism and redox biology in breast cancer and mammary epithelial cells which have undergone epithelial to mesenchymal transition

**DOI:** 10.1101/2025.04.08.647828

**Authors:** Haleigh N. Parker, Yongfeng Tao, Jenna Tobin, Kayla L. Haberman, Samantha Davis, Emily York, Alysia Martinez, Nobuyuki Matsumoto, Jaquelin Aroujo, Jun Hyoung Park, Bernd Zechmann, Benny Abraham Kaipparettu, Angela Boari, Christie M. Sayes, Antonio Evidente, Alexander Kornienko, Benjamin Cravatt, Daniel Romo, Joseph H. Taube

**Affiliations:** Department of Biology, Baylor University, TX, USA; Scripps Research Institute, La Jolla, CA, USA; Department of Chemistry and Biochemistry, Baylor University, Waco, TX, USA; Department of Molecular and Human Genetics, Baylor College of Medicine, TX, USA; Center for Microscopy and Imaging, Baylor University, TX, USA; Institute of Sciences and Food Production, National Research Council, Bari, Italy; Department of Environmental Science, Baylor University, Waco, TX, USA; Institute of Biomolecular Sciences, National Research Council, Pozzuoli, Italy; Department of Chemistry and Biochemistry, Texas State University, San Marcos, TX, USA

**Keywords:** Ophiobolin A, Epithelial-mesenchymal transition, Mitochondria, SLC25A40

## Abstract

Breast cancer progression is facilitated by the epithelial to mesenchymal transition (EMT), generating cancer cells with enhanced metastatic capacity and resistance to chemotherapeutics. EMT is known to impart changes in metabolic pathways and mitochondrial function. Here, we show a natural product possesses EMT-specific cytotoxic activity via alterations in metabolic and mitochondrial functions. The fungus-derived sesterterpenoid, ophiobolin A (OpA), possesses nanomolar cytotoxic activity and a high therapeutic index, although its target and mechanism of action remain unknown. Here, we utilized a model of mammary epithelial cells and breast cancer cell lines with and without EMT features to characterize the mechanism of selectivity towards EMT(+) cells by OpA and to identify novel targets for the treatment of EMT-enriched breast cancer. Proteins interacting with OpA in EMT(+) cells were identified through proteomic studies. We utilized trans-mitochondrial cybrids to determine that mitochondria mediate sensitivity to OpA. Furthermore, we report effects on glycolysis, oxidative metabolism, and disruption of metabolite abundance in the TCA cycle. Alterations to mitochondrial iron concentrations and glutathione production, mediated partly by OpA engagement with the metabolic proteins CISD3 and SLC25A40, were also detected in EMT(+) cells. Antioxidant mechanisms are activated by OpA in EMT(+) cells via the NRF2-ARE pathway, verified by decreased cytotoxicity in EMT(+) cells pretreated with the NRF2 activator CDDO. Collectively, we conclude that OpA selectivity toward EMT is mediated by the mitochondria, and at sub-cytotoxic levels, generates a metabolic shift leading to cell death countered by antioxidant mechanisms.

## Introduction

Breast cancer constitutes approximately 1 in 3 new cancer diagnoses in women, making it one of the leading female malignancies. It is estimated that 1 in 8 women will receive a breast cancer (BC) diagnosis in their lifetime (1). A heterogeneous disease, the treatment route and survivorship are often dictated by the molecular subtype of the cancer. Categorized by patterns in gene expression, BC can be described as luminal A, luminal B, HER2-enriched, basal-like, or claudin-low (2, 3). Via immunohistochemistry (IHC), BC can be further described by the presence of estrogen receptor (ER) and/or progesterone receptor (PR) constituting hormone receptor positive breast cancer (HR+), HER2-positive or triple negative breast cancer (TNBC) (2–4).

Negative for hormone receptors and HER2, TNBC accounts for 10-15% of all breast cancers and carries the lowest 5-year survival rate (4, 5). Most commonly diagnosed in women under 40, once TNBC has achieved distant metastasis, the 5-year survival rate drops precipitously to 12% (6). TNBC itself is heterogeneous and may be enriched for cells that have undergone epithelial to mesenchymal transition (EMT) (2, 7, 8). EMT is a conserved cellular program that causes epithelial cells to transition to a mesenchymal phenotype and gain adaptive characteristics such as enhanced motility, resistance to stress, and dedifferentiation potential (7, 9). In cancer, this program is hijacked, promoting drug resistance and metastasis. Initiated by extracellular cues such as growth factors, hypoxia, and inflammation, EMT is driven by master EMT transcription factors (EMT-TFs) *SNAI1* (Snail), *SNAI2* (Slug), *TWIST1* (TWIST) and zinc-finger E-box-binding homeobox 1 and 2 (ZEB1/ZEB2) which alter gene expression (7, 9, 10). Targeting and eliminating EMT(+) cells using drug-like small molecules has proven difficult (11–13), but this strategy holds great promise in reducing metastasis and chemotherapy resistance.

Possessing selective, anti-cancer cytotoxicity, which can be tuned to the acidic tumor microenvironment (14), the natural product ophiobolin A (OpA) has been investigated in the context of several cancers to determine the mechanism of action and selectivity (15–19). A sesterterpenoid derived from fungi in the genus *Bipolaris*, OpA targets metabolic processes in some cancer types (18, 20–22). The selectivity of OpA towards EMT(+) BC cells was demonstrated by increased sensitivity to OpA in EMT(+) cells in addition to decreased migration and colony-forming ability when cells were incubated with sub-cytotoxic concentrations (23). In conjunction with an increase in the expression CDH1 (E-cadherin) and a decrease in expression of CDH2 (N-cadherin), these results indicate a selective elimination of EMT(+) cells by OpA (23). However, the mechanism underlying this selectivity remains unknown. Recently, in lung cancer cells, OpA was observed to target two proteins in complex IV of the electron transport chain: HIGD2A and COX5A (22). Increases in oxidative metabolism, decreased membrane potential, and ATP production led the authors to conclude that OpA increases oxidative metabolism, leading to eventual metabolic collapse (22). These studies and others led us to test the impact of OpA on EMT-induced changes in metabolism.

To uncover the mechanism of OpA toward EMT in breast cancer, we examined the effect of OpA on ER+ MCF7, EMT(-), and TNBC MDA-MB-231, EMT(+), breast cancer cells as well as a direct model of EMT, comprising immortalized human mammary epithelial (HMLE) cells and HMLE cells expressing EMT-TF TWIST1 (HMLE-Twist). Utilizing mitochondrial cybrids and a panel of metabolic analyses, we report mitochondrial-mediated selectivity with specific effects on complex III of the electron transport chain and oxidative metabolism as a whole in EMT(+), but not EMT(-), breast cancer cells, generating a dependency on oxidative metabolism. Furthermore, we have identified several proteins as specific targets of OpA in EMT(+) TNBC cells, including CISD3 and SLC25A40, through proteomic studies. Engagement of these proteins by OpA was found to be associated with increased mitochondrial iron and glutathione production. Pointing to a ROS-mediated mechanism, we report activation of the NRF2 pathway, accumulation of cellular ROS and increased transcription of antioxidant associated genes in OpA-treated EMT(+) cells, with modulation of OpA cytotoxic activity via NRF2 activators or the antioxidant NAC. Our work indicates that OpA sensitivity is mediated via EMT-induced metabolic dependencies on ROS-limiting pathways, which may be overwhelmed at higher concentrations of OpA.

## Methods

### Cell Culture

MCF7 and MDA-MB-231 cell lines were purchased from ATCC; immortalized human mammary epithelial (HMLE) and HMLE-Twist cell lines were obtained from Dr. Sendurai Mani (Brown University). MCF7 and MDA-MB-231 cells were cultured in Dulbecco’s Modified Eagle’s Medium (DMEM) (Corning, Mediatech Inc., Manassas, VA, USA) with antibiotics (Penicillin/Streptomycin) (Gibco) and 10% fetal bovine serum (FBS) (Gibco, Fisher Scientific, Hampton, NH, USA). HMLE and HMLE-Twist cells were grown in a 1:1 ratio of Mammary Epithelium Basal Medium (MEBM) (Lonza, Walkersville, MD, USA) with Mammary Epithelial Growth Supplement (MEGS) (Gibco,) and Dulbecco’s Modified Eagle Medium (DMEM)/F12 1:1 (Cytiva, HyClone, Logan, UT, USA) with Penicillin/Streptomycin (Gibco), 5 μg/ml insulin (Sigma-Aldrich Co., St. Louis, MO, USA), 10 ng/ml human epidermal growth factor (EGF) (Millipore Corp., Billerica, MA, USA), and 0.5 μg/ml hydrocortisone (Acros, Fisher Scientific, Hampton, NH, USA) as in Yang et al. (24). All cell lines were grown and maintained at 37°C, 5% CO_2_, and consistently tested for mycoplasma.

### RT-qPCR

Cells were grown, as described above, until at least 85% confluency, then treated with diluted OpA or DMSO vehicle and incubated at 37C, 5% CO2. Cells were removed from plates and lysed using Trizol Reagent (Thermo Scientific, Waltham, MA, USA), and total RNA was extracted per manufacturer’s guidelines. Extracted RNA was quantified using a NanoDrop Microvolume Spectrophotometer (Thermo Scientific). cDNA was generated according to manufacturer recommendations. Comparative Ct method was used for relative mRNA quantification using the formula 2-*ΔΔ*Ct, and GAPDH was used for normalization. Primers for HO-1, xCT, GCLC, TRX, GPx1, CISD3, and SLC25A40 were obtained from Integrated DNA Technologies (Newark, NJ, USA). AzuraView GreenFast qPCR Blue Mix LR was obtained from Azura Genomics (Raynham, MA, USA).

### Seahorse Metabolic Assays

Cells were removed from culture plates with 0.15% trypsin (1:1 ratio of 0.05% trypsin (Corning,) and 0.25% trypsin (Gibco)) then seeded at 50,000 cells per well in a Seahorse XFp Cell Culture Miniplate (Agilent, Santa Clara, CA, USA) and incubated overnight for adherence. Cells were treated with either DMSO vehicle or diluted OpA. One day prior to use, a Seahorse XFp sensor cartridge was hydrated using Seahorse XF Calibrant Solution (Agilent) and placed in a non-CO2 incubator at 37°C overnight, according to manufacturer guidelines. On the day of testing, kit compounds were suspended and diluted in Seahorse XF assay media supplemented with 1mM pyruvate, 2mM glutamine and 10mM glucose, included with the kit, as per manufacturer instructions. Cell media was replaced with Seahorse XF assay media with supplements prior to performing each assay. Oxidative metabolism was determined using Agilent Seahorse XF Cell Mito Stress Test Kit. Glycolytic metabolism was analyzed using Agilent Seahorse XF Glycolytic Rate Assay Kit. ATP rate was measured using Agilent Seahorse XFp Real-Time ATP Rate Assay Kit.

### Transmission Electron Microscopy

Cells were fixed and stained as described by Parker et al. (25). In brief, cells were grown until at least 85% confluency after incubation with OpA or vehicle, washed twice with phosphate buffered saline (PBS), and then fixed for 30 min in 2.5% Glutaraldehyde dissolved in 0.06 M phosphate buffer. The samples were washed with PBS in triplicate for 10 min each before a secondary chemical fixation of 1% osmium tetroxide and 0.8% potassium ferrocyanide for 30 min was conducted. Cells were washed prior to serial dehydration with ethanol (EtOH) incubations for 10 min each step: one of 50%, one of 70%, two of 90%, and two of 100%.

Cells were infiltrated with Embed 812 to ethanol (1:2, 1:1, 2:1) then polymerized with an incubation at 60°C for 48 hrs. The samples were sectioned (80 nm) and post-stained with lead citrate and 1% uranyl acetate. The samples were imaged using transmission electron microscopy (TEM) (ThermoFisher Spectra 300 S/TEM). Mitochondrial area was analyzed using Olympus CellSens Dimension software (Olympus America Inc., Version 2.2). The ratio of roundness was determined as described by Lujan et al. 2021 (26). At least 8 cells were measured per cell type, with at least 50 mitochondria per cell line. Mitochondrial area, perimeter, elongation, and roundness were determined with subsequent analysis using GraphPad.

### Trans-Mitochondrial Cybrids

We have previously published the SUM159 cybrid models (27). The cybrids were generated using our previously published protocol (28). Briefly, mtDNA-depleted SUM159 cells (SUM159 ρ0 cells) were used as nuclear donors and were fused with enucleated A1N4 (benign) and SUM159 (TNBC) cells (mitochondrial donors) to generate A1N4/159 and 159/159 cybrids, respectively.

### Clonogenic assay

Clonogenic assays were performed as described previously (29). In a 6-well plate, 2,000 cells were seeded in triplicates. After 24 hrs, cells were treated with drugs and cultured for 10 days.

### Cell Viability in Cybrid Models

Cell viability assay was conducted using MTT (3-[4,5-dimethylthiazol-2-yl]-2,5 diphenyl tetrazolium bromide) colorimetric assay as previously published protocol (30).

### Electron Transport Chain (ETC) Complex Activity

Electron transport chain (ETC) enzymatic activity and citrate synthase (CS) activity assays were measured by kinetic spectrophotometric assays as previously described (31–33). The total protein amount of cell lysates was quantified by Bradford protein assay and used for normalization.

### Mitochondrial DNA (mtDNA) Copy Number Quantification

The quantitative RT-PCR of genomic DNA was performed by using the SYBR® Green PCR Master Mix with mtDNA and nuclear DNA specific primers (34). The copy number of mtDNA was calculated as described before (35).

### Ophiobolin A

OpA was produced by *Drechslera gigante*a and purified from the fungal culture filtrates, as reported previously (36). The fungus was grown and maintained on Petri dishes containing PDA (Oxoid, England). The culture filtrates were lyophilized, redissolved in distilled water, and extracted with EtOAc. The combined organic extracts were dehydrated, filtered, and evaporated under reduced pressure. The brown oily residue was fractionated by column chromatography, and crude OpA crystallized into white needles.

### Protein-directed ABPP

#### Cell treatment

MDA-MB-231 or HMLE-TWIST cells (20 million cells typically) were treated with competitors at indicated concentrations for the indicated time. Subsequently, the cells were treated with the alkyne probes at indicated concentrations for 1 h. Cells were then harvested and stored as pellets at -80 °C.

#### CuAAC conjugation of an azide-biotin tag to stereoprobe-labeled proteins

To the cell plate, 550 µL of cold DPBS was added, and cell lysis was achieved by sonication (2×8 pulses). Total protein quantification was performed using Pierce^TM^ BCA protein assay kit from Thermo Scientific as described below, and samples were normalized by dilution with PBS to a final concentration of 1,950 µg/mL in 500 µL. To the cell lysate, 55 µL of a labeling mixture (CuSO_4_: 10 µL of a 50 mM solution in MQ-water; Biotin-PEG3-N_3_: purchased from ChemPep Inc., cat #271605: 5 µL of a 10 mM DMSO stock, Tris(2-carboxyethyl)phosphine (TCEP), purchased from Biosynth®: 10 µL of a 13 mg/mL solution in MQ-water and Tris[(1-benzyl-1H-1,2,3-triazol-4-yl)methyl]amine (TBTA): 30 µL of a 1.7 mM solution in DMSO:tBuOH 1:4 v/v). Final concentrations for the copper-catalyzed azide-alkyne cycloaddition (CuAAC) reactions: 2 mg/mL proteome, 1 mM CuSO_4_, 0.1 mM Biotin-PEG3-N_3_, 1 mM TCEP and 0.1 mM TBTA. The mixture was incubated at r.t. for 1 hr and mixed vigorously every 15 min. 500 µL of MeOH and 100 µL of CHCl_3_ were added, the samples were vortexed and centrifuged at 16,000 × g at 4°C for 15 min. A protein disk was observed at the interphase, and the liquid layers were discarded. 1 mL of MeOH was added, the samples were mixed, centrifuged at 16,000 × g at 4°C for 15 min and the liquid layer was discarded. The solid protein palette was air-dried for 15 h and stored at -80 °C for further use.

#### Protein enrichment, tryptic digestion, and TMT labelling of tryptic peptides

To each sample, 500 µL of 8 M urea in DPBS and 10 µL of 10% of sodium dodecyl sulfate (SDS, purchased from BioPioneer Inc.) in MQ-water was added, and the samples were homogenized by sonication (8 pulses). 25 µL of a 200 mM DTT solution in MQ-water were added, and the samples were incubated at 65 °C for 15 min, before at r.t. 25 µL of a 400 mM iodoacetamide solution in MQ-water was added, followed by incubation in the dark at 37 °C for 15 min. Each sample was diluted by adding 5.5 mL of PBS and 130 µL of 10% SDS in PBS. 50 µL of streptavidin agarose beads (50% slurry) were washed with PBS (3 × 200 µL), resuspended in 100 µL PBS, and added to the samples. The samples were rotated at 37 °C for 2 hr. Then, the samples were centrifuged at 2,000 × g for 2 min (pelleted), and the liquid phase was discarded. The samples were washed by consequent addition/pelleting/liquid discard of 2 × 1 mL 0.2% SDS in PBS, followed by 1 × 1 mL, 2 × 1 mL MQ-water, and 1 × 1 mL 200 mM EPPS pH 8. The beads were then resuspended in a resuspension buffer (2 M urea, 200 mM EPPS, pH 8), and to each sample 4 µL of trypsin (1 µg modified trypsin, resuspended in 3 µL of trypsin resuspension buffer and 1 µL of 100 mM CaCl_2_) was added, and the samples were incubated under shaking at 37 °C for 15 h. The samples were resuspended and filtered over a a Biospin^TM^ column by centrifugation at 2,000 × g while collecting the flow through (ca. 200 µL). 85 µL of MeCN were added to achieve 285 µL of 30% MeCN aq. solution. 6 µL of the corresponding TMT tag (solution in MeCN; TMT10plex^TM^ isobaric label reagent purchased from ThermoFisher) were added, and the samples were mixed and incubated at r.t. for 75 min. Then, 6 µL of 5% hydroxylamine (purchased from Sigma Aldrich) were added, and the samples were incubated at r.t. for a further 15 min. Formic acid (10 µL) was added, and all samples of one 10-plex were combined into one Eppendorf tube and concentrated using the Speedvac SPD2030.

#### Peptide fractionation

The dry sample was resuspended in 0.5 mL of buffer A (95% H_2_O, 5% MeCN, 0.1% FA), and 40 µL of formic acid was added. The sample was sonicated in a water bath for 5 min. Acidic pH (pH < 3) was tested using pH paper. For de-salting the sample, a SepPak® C18 column from Waters, pre-conditioned with 100% MeCN and equilibrated with 3 × 1 mL buffer A was used. Then, the sample was added dropwise (1 drop/sec), and the flow-through was reloaded dropwise. The column was washed with 3 × 1 mL buffer A, and the peptides were eluted with 1 mL buffer B (80% MeCN, 20% H_2_O, 0.1% FA) and concentrated using the Speedvac SPD2030. The dry sample was resuspended in 0.5 mL of buffer A and sonicated in a water bath for 5 min. A preparative HPLC system was used for fractionation. The C18 column was conditioned with 80% MeCN, 20% aq. 10 mM (NH_4_)_2_CO_3_ for 15 min, then equilibrated in 100% aq. 10 mM (NH_4_)_2_CO_3_ and the sample was injected. A gradient from 100% aq. 10 mM (NH_4_)_2_CO_3_ to 80% MeCN, 20% aq. 10 mM (NH_4_)_2_CO_3_ was applied over 115 min total run time, and 96 fractions were collected in a 96-well plate pre-treated with 20 µL of 20% FA, 80% H_2_O in each well. The fractions on the plates were concentrated using the Speedvac SPD2030. Using a 12-chanel multipipette 100 µL of buffer B was added to the first row of the 96-well plate, and wells of one column (e.g. wells 1, 13, 25, 37, 49, 61, 73, 85) were subsequently washed. The procedure was repeated 3 times to combine the fractions in a final volume of 300 µL. The fractions were concentrated using the Speedvac SPD2030 and directly used for mass spectrometry analysis.

#### Filters for data processing

all channels normalized to a combined total value of 100. Control condition (only DMSO, no compound or alkyne) has a value <5; Standard deviation of each condition <5; significant competition, assigned as percentage competition of (active competitor, 100 nM) >=60. The data after filtering was used to generate the heatmap as Fig 1D.

**Figure 1.**
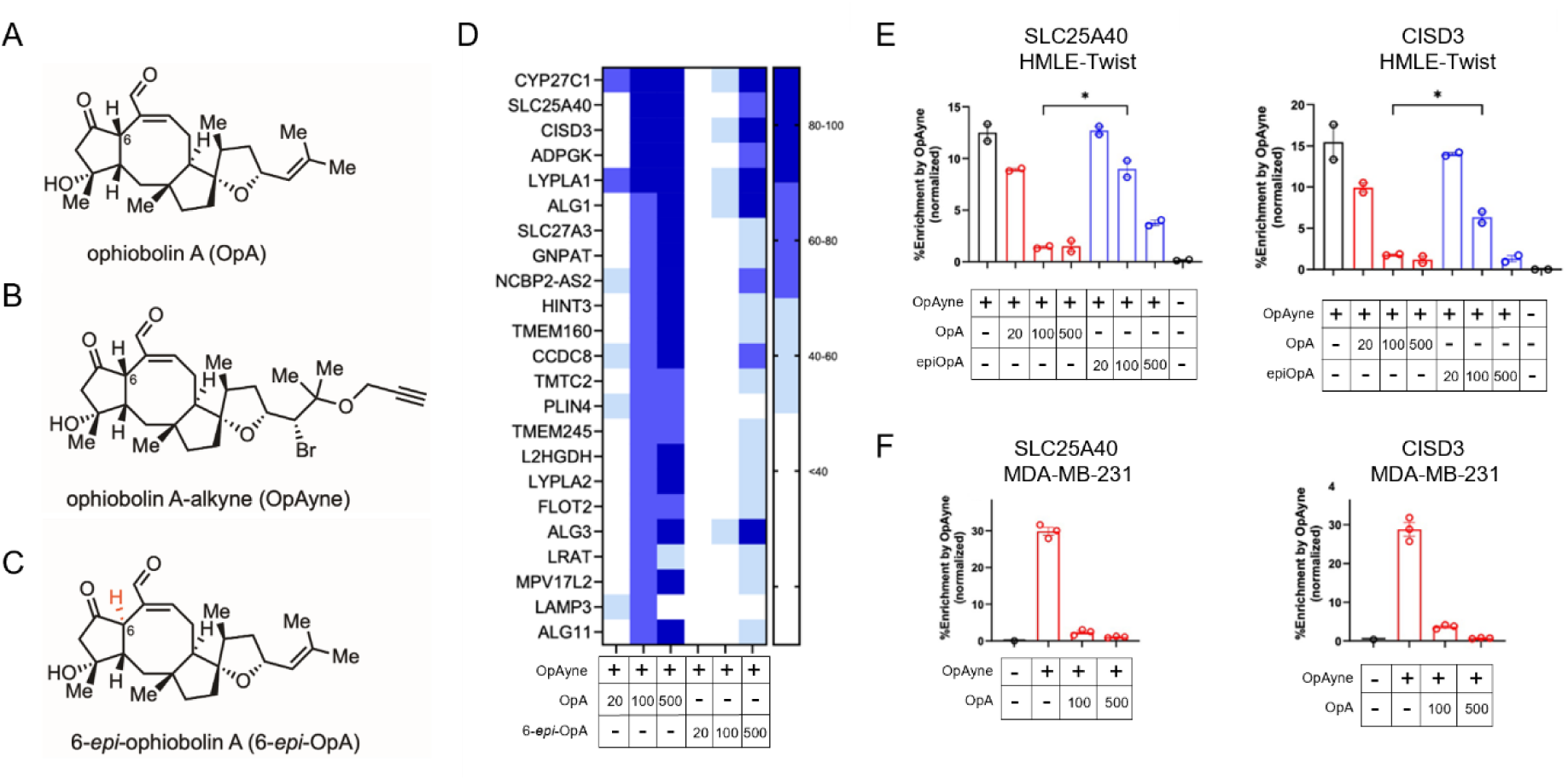
OpA Interacts with Metabolically Associated Proteins. Molecular structure of **A** ophiobolin A (OpA), **B** OpA-alkyne probe (OpAyne), and **C** 6-epi-ophiobolin A (6-epi-OpA). **D** Heatmap displaying the capacity of OpA or 6-epi-OpA to block protein engagement by the OpA-alkyne probe. **E-F** Bar-graphs displaying the percentage of protein engagement by the OpA-alkyne probe, with or without pre-treatment with OpA or epi-OpA at the indicated concentrations. n = 3 biological replicates.

### Targeted Metabolomics

Cells were grown and maintained to at least 80% confluency, as previously described. Once confluent, cells were treated with either DMSO vehicle, 50nM, or 125nM OpA (suspended in DMSO, diluted in PBS) for three hours at 37°C, 5% CO_2_. Cells were then pelleted and diluted to approximately 4 million cells per sample. Pre-analysis, cell pellets were snap-frozen in LN2. Liquid chromatography -mass spectrometry was performed on samples by Baylor College of Medicine Metabolomics Core.

### Cell Viability

Once confluent, cells were trypsinized and plated in 96-well plates at approximately 10,000 cells/well. Post overnight adherence, diluted OpA was added to each well and incubated for 24 hr at 37°C, 5% CO_2_. Cell Titer Blue (Promega, #G8080) was mixed into each well and incubated for four hours. Absorbance was read in on an AccuSkan Go Plate Reader (Fisher Scientific, Hampton, NH) (ex/em: 560/590).

### CDDO and NAC

Once confluent, cells were trypsinized and plated in 96-well plates at approximately 10,000 cells/well. After adhering overnight, cells were pretreated with either 10nM CDDO (MedChem Express, #HY-15725) or 10mM N-acetyl cysteine (NAC) (Sigma Aldrich, #A7250-10G) for 24 hours. CDDO was suspended in DMSO, diluted in PBS. NAC was suspended and diluted in molecular grade H2O. Media containing either vehicle, CDDO or NAC was removed and replaced with media containing various concentrations of OpA, and incubated for an additional 24 hours. Incubations were performed at 37C, 5% CO2. Cell Titer Blue (Promega, #G8080) was mixed into each well and incubated for four hours. Absorbance was read on an AccuSkan Go Plate Reader (Fisher Scientific, Hampton, NH) (ex/em: 560/590).

### Citrate Synthase

Cells were grown and maintained as described above, then treated with either vehicle, 50nM or 125nM OpA for three hours at 37C, 5% CO2. Cells then underwent subcellular fractionation, performed according to manufacturer guidelines in Subcellular Protein Fractionation Kit for Cultured Cells (Thermo Scientific, #78840). Protein content was determined in each sample via Pierce BCA Protein Assay Kit (Thermo Scientific, #23225), according to manufacturer guidelines. Equal amounts of protein were then used in the Citrate Synthase Activity Assay Kit (Abcam, #ab119692) to determine enzymatic activity. Object density as 412nm wavelength was measured on an AccuSkan Go Plate Reader (Fisher Scientific, Hampton, NH) every 5 min for 40 min. The change in activity over time was then determined.

### Glutamate Dehydrogenase

Activity was measured with the Glutamate Dehydrogenase Enzyme Activity Assay kit (ab102527, Abcam, Cambridge, UK) according to the manufacturer’s instructions. 10 x 10^6^ cells of each cell line were grown and treated with different concentrations of OpA treatment for 3 hr at 37°C, 5% CO_2_. Both treatment and control groups were lysed and then added to the 96-well microplate containing the reagents of the kit. The samples were mixed with the reaction reagents and then detected at 450 nm with the Thermo Scientific Multiskan Go at 450 nm. Glutamate Dehydrogenase activity was calculated after a 40-minute incubation and was repeated biologically 3 times.

### Flow Cytometry

Cells were grown and maintained as described above. Post-OpA treatment, cells were removed from plates and diluted to approximately 500,000 cells/mL. Cells were then incubated with 100nM MitoTracker Red CMXRos (Invitrogen, #M7512) for 30 min, washed with PBS, and incubated with 50 nM MitoTracker Green (Invitrogen, #M7514). Before final resuspension in FACs buffer (1% FBS in PBS), cells were incubated with one μM Sytox Blue viability stain (Invitrogen, #534857). Flow cytometry was performed on the FACS Melody (BD Biosciences, Franklin Lakes, NJ), and data was analyzed on FlowJo Analysis Software. The fluorescent intensity of MitoTracker Red and MitoTracker Green was determined post-exclusion of Sytox-positive cells.

### Immunoblotting

Protein was obtained from cells via RIPA Lysis buffer (Thermo, #J63306) containing phosphatase and protease inhibitors (Thermo, #78430) on ice. Protein content was quantified using Pierce BCA Protein Assay Kit (Thermo Scientific, #23225). Protein was run on 10% poly-acrylamide gel and transferred to PVDF membranes. Primary antibodies were diluted in 5% milk in TBST and incubated at 4C overnight with continuous rocking. Secondary antibodies were incubated for 1 hr at r.t.. Chemiluminescence was detected using ECL Prime (Cytiva, #RPN2236) and the Bio-Rad ChemiDoc system. Quantification of chemiluminescent images was performed on Gel Analyzer, with Actin as loading control. Primary antibodies include SLC25A40 (Cell Signaling, #E9C7Y), CISD3 (ProteinTech, #30480-1-AP) and Actin (Cell Signaling, #P6A8). The secondary antibody was Anti-Rabbit (Cell Signaling, #70745).

### Mitochondrial Iron

Mitochondrial iron accumulation post-treatment of OpA was evaluated using the iron-sensitive fluorescent probe RPA (rhodamine B-[(1,10-phenanthrolin-5-yl)-aminocarbonyl]benzyl ester) (BechChem, #B2852812). Cells adhered to the 96-well microplate after 24 hr in incubation and then were treated with varying concentrations of OpA for 3 hr at 37°C, 5% CO_2_. Cells were then incubated with HBSS and 1.5 μM RPA for 15 min at 37°C followed by two washes of HBSS. The average fluorescence was measured using the Bio Tek Lionheart FX Automated Microscope using cellular analysis.

### Glutathione

The concentration of free and oxidized glutathione was determined using the Glutathione Colorimetric Detection Kit (Invitrogen, #EIAGSCH), according to manufacturer guidelines. Briefly, cells were pelleted post OpA incubation and diluted to equal numbers of cells per sample. For free glutathione, cells were lysed, further diluted in assay buffer, plated in 96-well plates, and incubated with reaction buffer. Oxidized glutathione was determined by incubating samples with 2-vinyl pyrimidine (2VP, Acros Organics, #158002500) before dilution and plating. Post incubation with reaction buffer, fluorescent absorbance of samples and standards was read on an AccuSkan Go Plate Reader (Fisher Scientific, Hampton, NH) at 405nm. A standard curve was used to determine sample concentration. Values obtained from free and oxidized glutathione were used to determine total glutathione, per kit instructions.

### Cellular Reactive Oxygen Species (ROS)

Cellular ROS was evaluated in OpA treated cells using the DCFDA/H2DCFDA Cellular ROS Assay Kit (Abcam, #ab113851), according to kit instructions. Briefly, cells were plated on 96-well dark wall plates at approximately 25,000 cells/well. After adhering overnight, cell growth media was replaced with assay buffer containing DCFDA cellular ROS stain. After washing with assay buffer, cells were treated with OpA or vehicle, diluted in supplemental assay buffer for 3 hr. Fluorescence was measured on an AccuSkan Go Plate Reader (Fisher Scientific, Hampton, NH) (ex/em 485/535).

### NRF2 Nuclear Translocation

Cells were plated at approximately 25,000 cells on glass coverslips and allowed to adhere overnight in 6-well plates. Cells were then incubated with OpA for 3 hr at 37°C, 5% CO_2_, washed with PBS. Cells underwent fixation, first by incubating with 2% paraformaldehyde (Spectrum, #30525-89-4) in PBS. Cells were permeabilized by 0.3% Triton (Fisher Scientific, #BP151-100) in PBS and quenched with 1% glycine (Fisher Scientific, #BP381-5) in PBS. Cells were blocked with 8% BSA (Fisher Scientific, #BP1605-100) in TBST at 4°C overnight. The next day, coverslips were incubated with diluted NRF2 primary antibody (Cell Signaling, #D1Z9C) in a humidity chamber overnight at 4°C. Secondary anti-rabbit Alexa 488 antibody (Cell Signaling, #44125) was diluted in 3% BSA in TBST and incubated with cells for 1 hr at r.t. DAPI, diluted in H2O, was added to coverslips and incubated for 5 min at r.t.. Post wash, coverslips were mounted to slides using Permount (Fisher Scientific, #SP15-100) and sealed with clear nail polish. Slides were imaged using the Confocal Laser Scanning Microscope FV3000 (Olympus Corp., Tokyo, Japan), and images were analyzed using CellSens Imaging Software to determine the Pearson Correlation Coefficient.

### Generation of Knockdown Cell Lines

shRNA for scramble, CISD3 and SLC25A40 was obtained from Horizon Discovery. Bacterial growth for the generation of plasmids was performed using 100ug/mL Ampicillin (Fisher Scientific, #BP1760-25) resistant plates. Post colony growth, individual colonies were expanded in LB broth (Fisher Scientific, #BP1426-500). Plasmids from expanded bacterial colonies were isolated using the GeneJET Plasmid Midiprep Kit (Thermo Scientific, #K0481/0482), according to manufacturer guidelines. Lentiviral transduction was performed using 293T cells and FuGENE HD transfection reagent (Promega, #E2311). Protamine sulfate was added to the viral supernatant, filtered, administered to target cell lines and incubated for 24 hr. MDA-MB-231 knockdown cell lines were selected via puromycin. HMLE-Twist cell lines were sorted on the Sony SH800S Cell Sorter (Sony, New York, NY) for GFP positivity. Cell lines were frequently checked for GFP positivity and underwent selection/sorting as needed.

### Statistical Analysis

Statistical analysis was performed on GraphPad prism using 1way-ANOVAS, 2way-ANOVAS, false discovery rate (FDR), and t-tests with multiple comparison corrections. R studio was utilized to evaluate protein engagement datasets, using tidyverse, dplyr, tidyr and ggplot2 packages on R Markdown.

## Results

### OpA Interacts with Metabolism-Regulating Proteins

To begin elucidating the mechanism underlying OpA cytotoxic activity, we first investigated possible molecular targets. As OpA (Fig. 1A) contains a ketoaldehyde moiety, it is likely that the small molecule reacts with lysines or cysteines within target proteins as previously proposed (16, 37). In order to determine these possible targets, we conducted protein-directed activity-based protein profiling (ABPP) experiments, as described by Njomen, et. al. (38), by treating cells with the OpA-alkyne probe (OpAyne, Fig. 1B) to enrich interacting proteins. As a control, we pre-treated the cells with increasing concentrations of OpA or the phenotypically inactive congener 6-*epi*-ophiobolin A (6-*epi*-OpA (37), Fig. 1C) as competitors before adding the OpA-alkyne probe. The pre-treatment with OpA, but not 6-*epi*-OpA, would compete for phenotypically relevant binding sites, and thus reduction in protein enrichment by OpA, but not 6-*epi*-OpA pre-treatment, would strengthen the confidence in the specificity of the liganding event.

In HMLE cells with constitutive expression of the EMT-TF, Twist1 (HMLE-Twist), several proteins exhibited strong enrichment by the OpA-alkyne probe, with more than 60% competition observed when cells were pre-treated with 100 nM OpA but significantly less competition by pre-treatment of 6-*epi*-OpA at the same concentration (Fig. 1D, E). Examples include CISD3 and SLC25A40, both with metabolic or mitochondrial functions. In MDA-MB-231 cells, representative of an EMT(+) breast cancer cell line, both proteins also show substantial engagement by the OpA-alkyne probe that is blocked by pre-treatment with OpA (Fig. 1F).

### Sensitivity to OpA Induced Cell Death is Mediated by the Mitochondria

Based on OpA engagement with proteins associated with metabolism, we hypothesized OpA sensitivity may be linked to mitochondria. Thus, we utilized trans-mitochondrial cybrids to determine whether sensitivity to OpA is linked to mitochondrial or nuclear origin (Fig. 2) (39). EMT(+) SUM159 cells were selected for the nuclear background and non-transformed mammary epithelial A1 cells were selected as mitochondrial donors.

**Figure 2.**
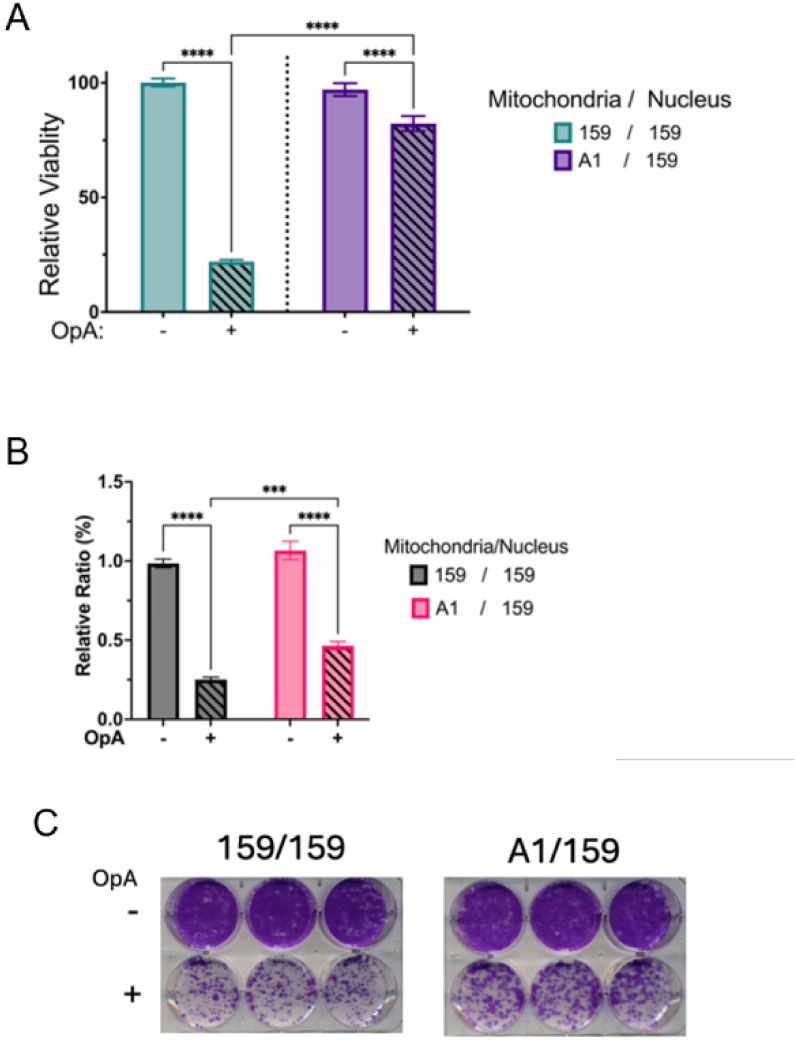
Sensitivity to OpA Induced Cell Death is Mediated by the Mitochondria. **A** Relative viability of cybrid cells post 72h incubation with either vehicle or 500nM OpA. **B,C** Relative ratio of colony growth in cybrid cells post 10 days incubation with either vehicle or 500nM OpA. Statistical analysis performed on GraphPad Prism using 2way-ANOVA and Šidák’s multiple comparison test or unpaired t test (*p<0.5; **p<0.01; ***p<0.001; **\*\*\*\***p<0.0001).

After incubation with OpA, the cybrid models were evaluated for changes in cell viability (Fig. 2A). In the SUM159 cells with A1 mitochondria (A1/159 cybrid), the sensitivity to OpA was significantly reduced relative to the 159/159 cybrid, indicating that A1-derived mitochondria confer a protective effect against OpA-induced cytotoxicity. The A1/159 cybrid model was further evaluated for the effect of OpA on colony formation. Incubation with OpA causes decreased colony formation in the 159/159 cybrid, an effect that is abrogated in the A1/159 cybrid (Fig. 2B/C). These responses to OpA indicate that sensitivity is dictated by mitochondria.

### OpA Induces Loss of Membrane Potential and Functional Mitochondria

We next sought to determine if OpA induces a decline in mitochondrial function by assessing mitochondrial membrane potential (MMP) through staining with MitoTracker Red. Consistent with the role of mitochondria in sensitivity to OpA, we observed a significant decrease in MMP in MDA-MB-231 cells treated with OpA at 50 nM and at 125 nM (Fig. 3A, left). However, in MCF7 cells, which exhibit lower overall MMP, this parameter was only decreased at 125 nM OpA, not at 50 nM (Fig. 3A, right). Notably, OpA reduces the MMP of MDA-MB-231 cells to approximately the same value as MCF7 cells. Similarly, treatment with 50 and 125 nM OpA significantly decreases MMP in HMLE-Twist cells (Fig. 3B, left) but not HMLE-vector cells (Fig. 3B, right). Similar to losses in MMP, we detected a significant decrease in the fluorescent signal of total active mitochondria, as measured by MitoTracker Green (Fig. 3C, D). While treatment with 50 and 12

**Figure 3.**
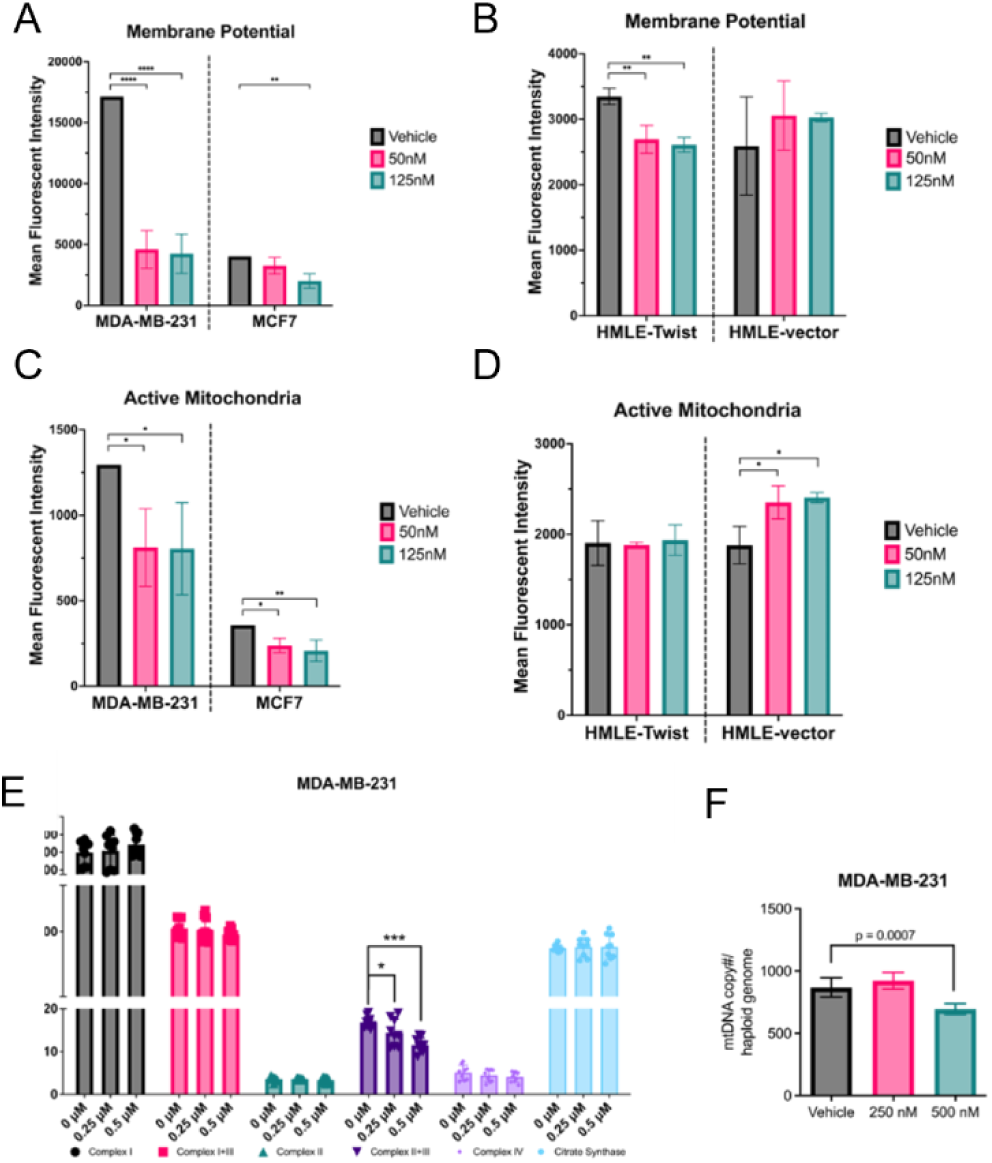
OpA Induces Loss of Mitochondrial Membrane Potential. Functional Mitochondria Mean fluorescent intensity of MitoTracker Red, representative of MMP, in **A** MDA-MB-231 and MCF7 cells and **B** HMLE-Twist and HMLE-vector cells. Mean fluorescent intensity of MitoTracker Green, representative of active mitochondria, in **C** MDA-MB-231 and MCF7 cells and **D** HMLE-Twist and HMLE-vector cells. Cells in A-D treated with either vehicle, 50nM or 125nM OpA for 3h. n = 3 biological replicates. **E** Activity of complexes in the ETC in MDA-MB-231 cells (average of three independent experiments). **F** Ratio of mtDNA copy number to haploid genome in MDA-MB-231 cells (average of two independent experiments). Cells in E, F treated with either vehicle, 250nM or 500nM OpA for 72h. Statistical analysis performed on GraphPad Prism using 1way-ANOVA and Dunnett’s multiple comparison test or unpaired t test (*p<0.5; **p<0.01; ***p<0.001; ****p<0.0001).

5 nM OpA significantly reduces total active mitochondria in both MDA-MB-231 and MCF7 cells, the magnitude of the change in greater for MDA-MB-231 cells (Fig. 3C). OpA does not cause significant changes to the level of total active mitochondria in HMLE-Twist cells (Fig. 3D).

To further describe OpA induced alterations to MMP, we analyzed changes in the activity of the complexes that make up the electron transport chain (ETC) in MDA-MB-231 cells, as these cells had the most significant response. Using 250 or 500 nM OpA, we found a dose-dependent decrease in the activity of complexes II and III (Fig. 3E). We also found a significant decrease in mtDNA copy number in MDA-MB-231 cells (Fig. 3F), supporting the previous observation of decreased total active mitochondria (Fig. 3C).

### OpA Alters Metabolite Abundance in the TCA Cycle and Glycolysis

Alterations to MMP can affect several metabolic pathways, including glycolysis and the TCA, which may then affect overall mitochondrial functionality (40, 41). Furthermore, as metabolic pathways work in tandem, significant changes to one pathway can cause dependency or collapse of another (41–43). Therefore, to assess OpA-induced changes to these pathways, we performed targeted metabolomics on MDA-MB-231 and MCF7 cells treated with 50 and 125 nM OpA for 3 hr (Sup. Fig. 1A, B).

At just 50 nM, OpA alters the abundance of 11 metabolites involved in the TCA cycle and glycolysis in MDA-MB-231 cells (Sup. Fig. 1A). Notably, pyruvate, fumarate, malate, and oxaloacetate are all increased. In comparison to the MDA-MB-231 cells, far fewer metabolites are significantly affected in the MCF7 cells, indicating a lesser effect on these pathways by OpA (Sup. Fig. 1B). Because oxaloacetate is increased and aconitate is decreased, we hypothesized OpA may decrease the activity of citrate synthase (CS), the enzyme which converts acetyl-CoA and oxaloacetate into citrate, which in turn becomes aconitate. In both MDA-MB-231 and MCF7 cells, neither 50 nM nor 125 nM OpA had a significant impact on CS activity (Sup. Fig. 1C). Similarly, as glutamic acid and ketoglutarate are also found to be increased in MDA-MB-231 cells, we measured the activity of glutamate dehydrogenase (GDH). As with CS, neither dose tested caused a significant alteration to GDH activity in either cell line (Sup. Fig. 1D). These results indicate that while OpA does not directly affect CS or GDH enzyme activity, metabolite abundance is significantly altered, possibly through alterations to the activity of the pathways as a whole.

### OpA Increases Oxidative Metabolism While Decreasing Glycolysis and ATP Production

Incubation with OpA at 50 nM causes dysregulation of several metabolites, yet did not alter the activity of key enzymes required for the synthesis of these metabolites (Sup. Fig. 1). We hypothesized this may be due to alteration of pathway activity and capacity as a whole, rather than at the enzymatic level. Moreover, EMT(+) cancer and non-cancer cells may dynamically shift between oxidative and glycolytic metabolic phenotypes to respond to energy demands. We further hypothesized that OpA alters this ability in EMT(+) cells by decreasing the capacity of pathways to meet energy needs. Thus, we sought to determine changes in the activity and plasticity of oxidative metabolism and glycolysis.

The oxygen consumption rate (OCR) was evaluated in the presence of several mitochondrial inhibitors to determine whether OpA induces changes in oxidative metabolism at 50 and 125 nM via the Seahorse Mito Stress Test. Basal respiration is not significantly affected in either cancer cell line, or the HMLE-vector cells (Fig. 4A, B). However, the HMLE-Twist cells have a slightly elevated basal OCR at 125 nM OpA (Fig. 4B). Measured after the addition of oligomycin and FCCP, maximal respiration is significantly increased in MDA-MB-231 cells, which was not observed in MCF7 cells (Fig. 4A). Similarly, maximal respiration is also increased in HMLE-Twist cells yet decreased in HMLE-vector cells (Fig. 4B). Lastly, the ability of each cell line to respond to an energy demand, or spare respiratory capacity, was evaluated after addition of all inhibitors. Comparable to maximal respiration, MDA-MB-231, and HMLE-Twist cells have significantly increased spare respiratory capacity (Fig. 4A, B). No significant change is observed in MCF7 cells, while spare capacity is decreased in HMLE-vector cells (Fig. 4A, B).

**Figure 4.**
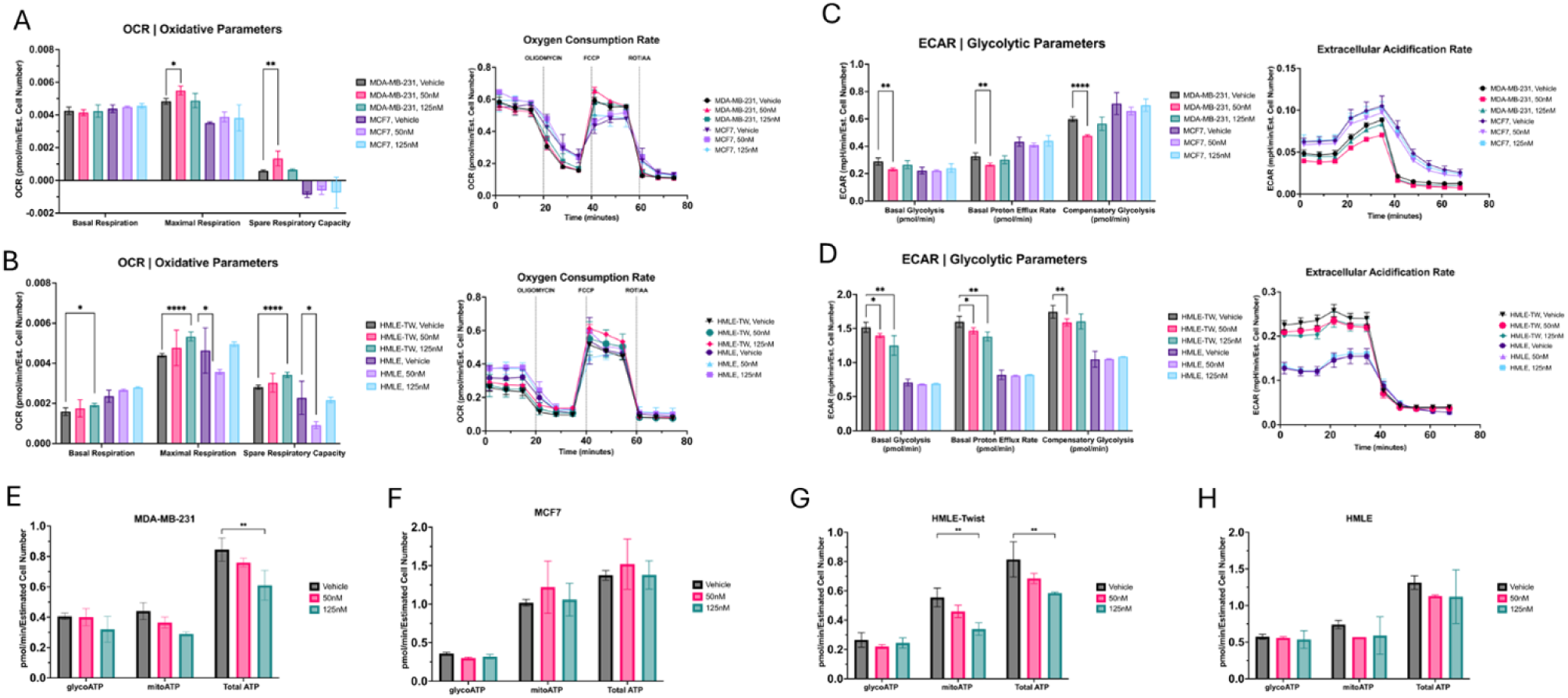
OpA Increases Oxidative Metabolism, while Decreasing Glycolysis and ATP Production. Parameters of oxidative metabolism in **A** MDA-MB-231 and MCF7 cells (OCR, right) and **B** HMLE-Twist and HMLE-vector cells. Representative seahorse plots included. Parameters of glycolytic metabolism in **C** MDA-MB-231 and MCF7 cells (ECAR, right) and **D** HMLE-Twist and HMLE-vector cells (ECAR, right). Representative seahorse plots included. E-H ATP production rate via glycolytic and mitochondrial metabolism in **E** MDA-MB-231, **F** MCF7, **G** HMLE-Twist and **H** HMLE-vector cells. All cell lines treated with either vehicle, 50nM or 125nM OpA for 3h. n = 3. Statistical analysis performed on GraphPad Prism using multiple unpaired t tests and Holm-Šidák multiple comparison test or 2way-ANOVA with Dunnett’s multiple comparison test (*p<0.05; **p<0.01; ***p<0.001; ****p<0.0001).

We next utilized the Seahorse Glycolytic Rate test to assess if any changes similar to those observed in oxidative metabolism were also present in glycolytic metabolism. As cancer cells are known to favor glycolysis, even in an aerobic environment, we hypothesized that OpA may have detrimental effects to this pathway, possibly contributing to the observed increase in oxidative metabolism. In both EMT(+) cell lines, basal glycolysis is significantly decreased. Neither EMT(-) cell line demonstrated significant changes (Fig. 4C, D) in the presence of 50 or 125 nM OpA. Correlated with lactate efflux and excluding CO_2_ dependent acidification, the basal proton efflux rate (PER) is also decreased in both EMT(+) cell line, but not the EMT(-) cell lines (Fig. 4C, D). Finally, after the addition of all inhibitors that preclude oxidative phosphorylation, compensatory mechanisms are forced to utilize glycolysis, allowing the capacity of glycolysis to compensate for energy needs to be measured. Importantly, compensatory glycolysis is also decreased in both EMT(+) cell lines, while neither EMT(-) cell line is significantly altered (Fig. 4C, D). This, in tandem with the observed increase in spare respiratory capacity (Fig. 4A,B), supports our hypothesis that OpA dysregulates metabolic plasticity by altering pathway activity and capacity.

The main product of both oxidative and glycolytic metabolism is ATP (44, 45). As both pathways are affected by OpA, we next sought to determine if ATP production is comparably affected. By measuring both OCR and ECAR in the presence of oligomycin and rotenone with antimycin A, ATP generated from either glycolysis (glycoATP) or the mitochondria (mitoATP) can be determined by the Seahorse ATP Rate test. In MDA-MB-231 and HMLE-Twist cells, OpA causes a dose-dependent decrease in total ATP production (Fig. 4E, G). No significant changes are observed at either concentration of OpA in the EMT(-) cell lines (Fig. 4F, H).

### OpA Elongates Mitochondria

Mitochondrial morphology is closely tied to its functionality. Availability of metabolites and oxygen, energy demands, and cellular movement can significantly affect mitochondrial shape and structure (46–49). As OpA shifts the activity of oxidative and glycolytic metabolism, we hypothesized this change would also be evident is mitochondrial morphology. Hence, we generated high resolution images using transmission electron microscopy (TEM) of each cell line treated with either OpA, at 50 or 125 nM or vehicle (Fig. 5).

**Figure 5.**
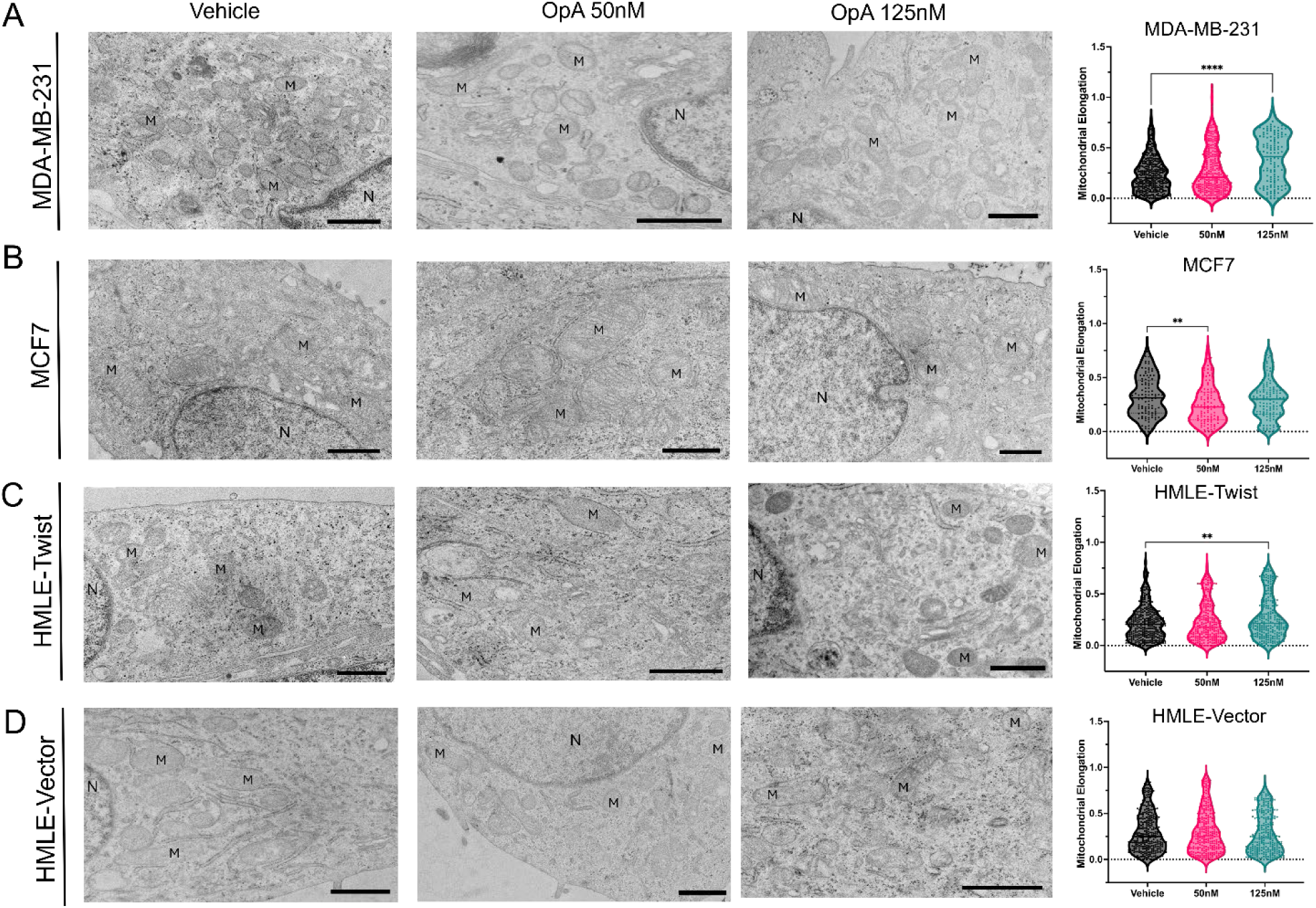
Mitochondria are Elongated Post OpA Treatment. TEM representative images (left) and quantification (right) of mitochondrial elongation in **A** MDA-MB-231, **B** MCF7, **C** HMLE-Twist and **D** HMLE-vector cells. N indicates the nucleus and M indicates mitochondria. Scale bar represents 1µm. n ≥ 105 mitochondria. Statistical analysis performed on GraphPad Prism using 1way-ANOVA with Dunnett’s multiple comparison test (*p<0.5; **p<0.01; ***p<0.001; ****p<0.0001).

Measuring individual mitochondria in MDA-MB-231 cells reveals an increase in elongation upon treatment with 125 nM OpA, with similar results in HMLE-Twist cells (Fig. 5A, C). However, mitochondria in MCF7 cells are less elongated when incubated with 50 nM OpA, while mitochondria in HMLE-vector cells are not significantly changed (Fig. 5B, D). These results support our hypothesis, indicating OpA affects the mitochondria, in EMT(+) cells, at both the functional and morphological level.

### OpA Disrupts Iron and Glutathione Homeostasis in EMT(+) Cells

We next evaluated the impact of OpA on molecular pathways downstream of CISD3 and SLC25A40. CISD3 is a mitochondrial transporter protein involved in maintaining the labile pool of mitochondrial iron and the regulation of ROS (50, 51). Therefore, we hypothesized OpA incubation would cause disrupted iron homeostasis in EMT(+) cells. Also a transporter protein, SLC25A40 is involved in glutathione (GSH) transport into the mitochondria (52, 53). Similarly, we hypothesized that OpA causes an alteration in GSH homeostasis in EMT(+) cells. Moreover, GSH is involved in the stability of iron/sulfur clusters, connecting the function of the two proteins (54–56).

To investigate changes to mitochondrial iron, we utilized fluorescent microscopy (Fig. 6A-D). As expected, no significant changes to iron accumulation in the mitochondria are detected in either EMT(-) cell line (Fig. 6B, D). However, contrary to our expectations, neither EMT(+) cell line exhibited a strong response, with statistical significance detected for HMLE-Twist cells, but not MDA-MB-231 cells, at 125 and 250 nM OpA (Fig. 6A, C). These results indicate that while CISD3 is an authentic target of OpA, this interaction does not necessarily lead to a direct functional outcome.

**Figure 6.**
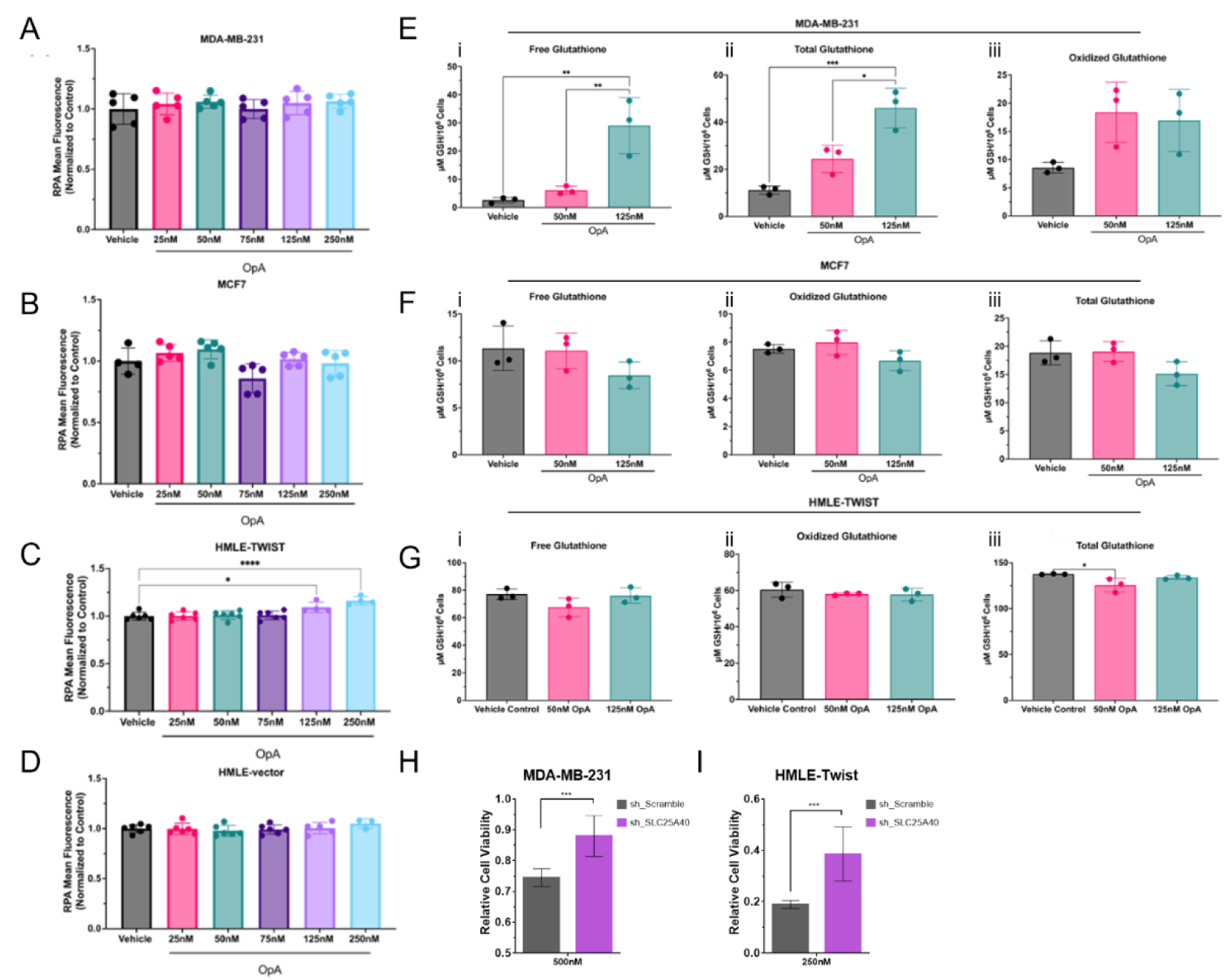
OpA Disrupts Glutathione Homeostasis in EMT(+) Cells and Requires SLC25A40 for cytotoxic activity. Relative fluorescence of mitochondrial iron marker RPA in **A** MDA-MB-231, **B** MCF7, **C** HMLE-Twist and **D** HMLE-vector cells treated with a range of OpA concentrations for 3h. n ≥ 4 replicates. Glutathione concentration per 10^6^ cells in **E** MDA-MB-231 cells, **F** HMLE-Twist cells and **G** MCF7 cells treated with either vehicle, 50nM or 125nM OpA for 3h. n = 4 replicates. Statistical analysis performed on GraphPad Prism using 1way-ANOVA with Tukey’s multiple comparison test. Relative viability of **H** MDA-MB-231 or **I** HMLE-Twist SLC25A40 knockdown or control cells treated with OpA for 24 hours. n = 6. Statistical analysis by Student’s t-test (*p<0.05; **p<0.01; ***p<0.001; ****p<0.0001).

Next, the cellular concentration of GSH and its oxidized form, GSSG, was ascertained in OpA treated cells (Fig. 6E-G). Free GSH is found to be dose-dependently increased in only the MDA-MB-231 cells (Fig. 6Ei), with no significant changes detected in HMLE-Twist, nor MCF7 cells (Fig. 6Fi, Gi). Interestingly, GSSG is not significantly changed in any cell lines, with MDA-MB-231 cells experiencing only about a two-fold increase with either OpA concentration (Fig. 6E-Gii). Total GSH is also largely unchanged in HMLE-Twist and MCF7 cells, with the latter demonstrating a slight decrease at 50nM OpA (Fig. 6Fiii, Giii). Importantly, total GSH is also dose-dependently increased in MDA-MB-231 cells, similarly to free GSH (Fig. 6Eiii). Therefore, the increases observed in total GSH are driven by increases in free GSH in OpA treated MDA-MB-231 cells. This may be due to decreased transport of GSH via SLC25A40, potentially disrupting its role as an antioxidant.

To ascertain whether expression of SLC25A40 affects OpA activity, we generated cells with reduced expression via shRNA-mediated knockdown (Sup Fig 2). We then sought to determine if decreased expression of SLC25A40 affects the cytotoxicity of OpA. OpA activity was abrogated in both MDA-MB-231 (Fig. 6H) and HMLE-Twist (Fig. 6I) cells upon SLC25A40 knockdown.

### OpA Activates NRF2-Mediated Antioxidation in EMT(+) Cells, Affecting Cytotoxicity

GSH is a potent antioxidant, upregulated during oxidative stress, with secondary functions in protein stability, cysteine storage and drug resistance (55, 57–59). Upstream of GSH, the NRF2 pathway can stimulate the production of GSH and other antioxidants via the activation of the antioxidant response element (ARE) (57, 59–61). As increased free and total GSH was observed in MDA-MB-231 cells, we next tested the hypothesis that OpA activates the NRF2-antioxidation response.

A set of transcripts is associated with the activation of the ARE, including heme oxygenase-1 (HO-1) and the cystine/glutamate antiporter xCT. While HO-1 is also associated with general oxidative stress, xCT is mainly associated with GSH production (59, 62, 63). Therefore, we evaluated changes in the expression of these genes in addition to others also associated with the ARE (Fig. 7A-D). In MDA-MB-231 cells, OpA stimulates an increase in transcription of HO-1 and xCT (Fig. 7A), while no significant changes are observed in MCF7 cells (Fig. 7B). OpA also causes a significant increase in HO-1 in HMLE-Twist cells (Fig. 7C) and xCT in HMLE-vector cells (Fig. 7D). We next quantified NRF2 translocation to the nucleus in EMT(+) cells via immunofluorescence. Upon treatment with 50 or 125 nM OpA, NRF2 translocation is significantly increased in MDA-MB-231 cells (Fig. 7E). Only at the higher concentration of OpA was increased translocation observed in HMLE-Twist cells (Fig. 7F). To further validate the activity of antioxidant mechanisms, we also measured cellular accumulation of reactive oxygen species (ROS). In both MDA-MB-231 and HMLE-Twist, a dose-dependent decrease is observed in cellular ROS, indicating antioxidation is occurring (Fig. 7G).

**Figure 7.**
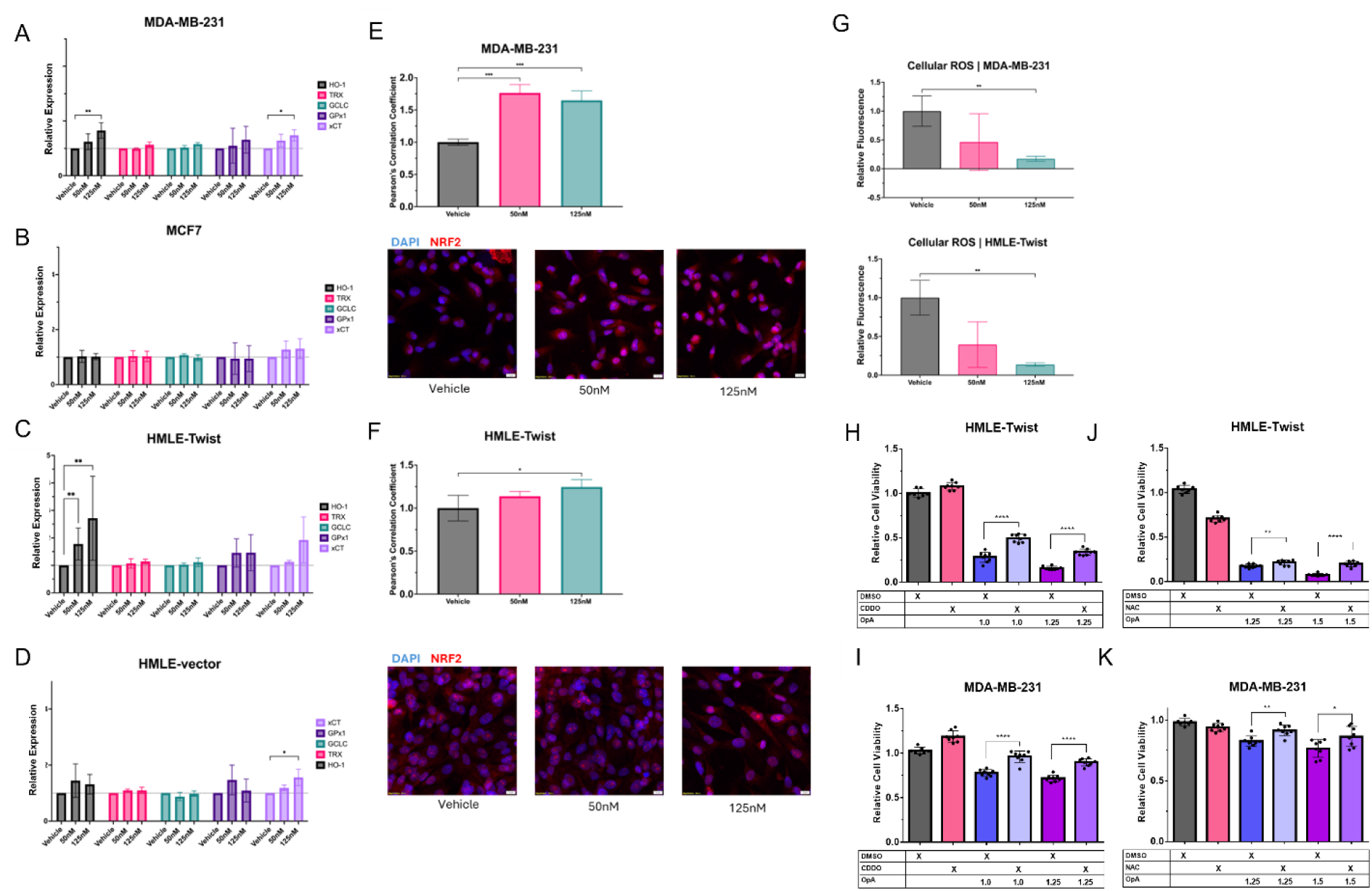
OpA Activates NRF2-Mediated Antioxidation in EMT+ Cells, Affecting Cytotoxicity. Transcriptional expression of genes associated with ARE activation in **A** MDA-MB-231, **B** MCF7, **C** HMLE-Twist and **D** HMLE-vector cells. n = 3 biological replicates. **E-F** Quantification (top) and representative images (bottom) of NRF2 nuclear translocation in **E** MDA-MB-231 cells and **F** HMLE-Twist cells. n = 3 biological replicates. **G** Relative fluorescence of cellular ROS marker DCFDA in MDA-MB-231 (top) and HMLE-Twist cells (bottom). n ≥ 4 replicates. **H-I** Relative cellular viability of MDA-MB-231 H and HMLE-Twist cells I pretreated with 10nM CDDO for 24 hours, then incubated with indicated concentrations of OpA for 24h. **J-K** Relative cellular viability of MDA-MB-231 J and HMLE-Twist cells K pretreated with 10mM NAC for 24 hours, then incubated with indicated concentrations of OpA for 24h. n = 8. Statistical analysis performed on GraphPad Prism using multiple unpaired t tests with Holm-Šidák multiple comparison test or 1way-ANOVA of Pearson’s Correlation Coefficient with Dunnett’s multiple comparison test (*p<0.05; **p<0.01; \***\*\***p<0.001; ****p<0.0001).

Next, to ascertain the role of the NRF2 pathway, MDA-MB-231 and HMLE-Twist cells were pretreated with the NRF2 activator CDDO to test the hypothesis that overactivation of the NRF2 pathway decreases OpA mediated cytotoxicity (59, 64). After a 24-hour incubation with high concentrations of OpA, cell death in MDA-MB-231 cells is partially rescued when pretreated with 10nM CDDO (Fig. 7H). Similar results are obtained in HMLE-Twist cells, with cell death significantly reduced at all concentrations of OpA with CDDO pretreatment (Fig. 7I). Cells were then pretreated with the antioxidant NAC before OpA incubation as another means of validating the involvement of antioxidant machinery. Again, cell death is significantly reduced at all concentrations of OpA with NAC pretreatment in MDA-MB-231 cells (Fig. 7J). HMLE-Twist cells are slightly more sensitive to NAC pretreatment. However, cell death was still significantly decreased at 1.5 μM and 1.25 μM OpA (Fig. 7J). Thus, we can conclude that OpA stimulates the NRF2 mediated antioxidant response in EMT(+) cells, abrogating cell death until these compensatory mechanisms fail and collapse.

## Discussion

The rewiring of cellular metabolic programming to meet the energetic demands of cancer cells introduces weaknesses which may be exploitable by therapeutic agents (65, 66). Natural products have historically been a source of novel investigative avenues for targeting these limitations and have led to effective, targeted cancer therapies. Here, we used models of EMT consisting of breast cancer and mammary cell lines enriched for or lacking EMT characteristics to determine the mechanism of EMT selectivity of the natural product OpA.

One Achilles heel that may enable the targeting of EMT(+) cells is the capacity of EMT to rewire cellular metabolism and mitochondrial function, increasing cellular metabolic plasticity, which is critical for enduring the metastatic cascade and metabolic insults received from anti-cancer agents (44, 67, 68). Cells generate ATP via oxidative phosphorylation (OXPHOS), glycolysis and fatty acid oxidation, with other pathways providing intermediates and metabolites (44, 45). In cancer cells, however, metabolism is reprogrammed towards glycolysis to meet the needs of uncontrolled cellular growth. Termed the Warburg effect, cancer cells will undergo aerobic glycolysis, quickly generating ATP and high levels of reactive oxygen species (ROS) (44, 45, 68). These alterations enhance metabolic plasticity, the ability of cells to respond to sudden metabolic demands and insults (69, 70). The necessity of metabolic plasticity in metastasis has been exemplified in experiments utilizing trans-mitochondrial cybrids. In cancer cells with mitochondria from a non-cancer cell line, invasion and colony formation, characteristics associated with EMT, were all significantly decreased, indicating decreased metastatic potential (39).

Utilizing an alkyne probe and mass spectrometry (Fig. 1), our data indicate that OpA, at 50 and 125 nM concentrations, engages with several metabolic regulators in EMT(+) cells. Therefore, we postulate that OpA’s mechanism of action is metabolically linked. Using trans-mitochondrial cybrids to investigate this theory, we determined that the level of OpA sensitivity is conferred by the mitochondria rather than the nucleus (Fig. 2). Using a panel of metabolic assays, we further described specific suppression of glycolytic capacity in EMT(+) cells to a level commensurate with EMT(-) cells, forcing cells to upregulate OXPHOS and elongate mitochondria (Fig. 3-5). Nevertheless, such adaptations fail to maintain ATP, paving the way for metabolic collapse and cell death.

Metabolic reprogramming in cancer cells is also associated with increased accumulation of damaging byproducts, such as ROS (39, 71), which elicit the NRF2-mediated antioxidant response (60, 61). Bound in the cytoplasm by KEAP1, NRF2 is released and translocates to the nucleus upon an accumulation of ROS (57, 59, 61). Then, acting as a transcription factor, NRF2 binds antioxidant response elements (ARE), activating the expression of antioxidant enzymes such as heme-oxygenase 1 (HO-1) and glutamate-cysteine ligase (GCL), the rate-limiting and initial enzyme of glutathione (GSH) synthesis (59–61). GSH, itself, is a primary cellular antioxidant and is commonly increased in cancer cells (55, 57). GSSG, or oxidized GSH, is readily reduced by NADPH and glutathione reductase (GR) to maintain a high ratio of GSH:GSSG (55, 57, 72). However, reliance on these pathways due to EMT-induced metabolic reprogramming introduces new weaknesses and targets to be exploited by novel therapeutic approaches (72, 73).

By interrogating OpA-interacting proteins, we identified that SLC25A40, transporter of GSH, is necessary OpA activity (Fig. 6). Furthermore, we link the NRF2-mediated antioxidant response to decreased OpA induced cytotoxicity by pretreating EMT(+) cells with either an NRF2 activator or ROS scavenger (Fig. 7). Collectively, our data illustrate a mechanism whereby OpA compromises mitochondrial function via disruption of mitochondrial GSH transport, pushing cells to an oxidative phenotype. This results in the activation of antioxidant machinery in EMT(+) cells, initially decreasing ROS, which may eventually be overwhelmed due to alterations in oxidative and glycolytic metabolism.

The mechanism of action employed by OpA has been investigated in diverse contexts. Using a panel of cancer cell lines, OpA was observed to induce context-dependent effects based on the cancer type (18). Interestingly, this study notes decreases in mitochondrial area in most cell lines and decreased membrane potential in several cell lines, including the TNBC cell line MDA-MB-231, when treated with OpA (18).

The mechanism by which mitochondria confer OpA sensitivity at low concentrations (50 nM) is likely linked to the crosstalk occurring between the mitochondria and nucleus during EMT (39, 74). Previous studies using similar trans-mitochondrial cybrids reported characteristics associated with EMT and metastasis, like invasion and migration, to be conferred by mitochondrial crosstalk with the nucleus (39). Additionally, mitochondrial retrograde signaling has been observed to induce EMT in mammary cells (74). Therefore, OpA targets may be upregulated after metabolic reprogramming via EMT-associated mitochondrial crosstalk (39, 74). Furthermore, OpA has also been reported to reverse EMT phenotypes, including the expression of E cadherin and N cadherin and cancer stem-like cell markers CD44 and CD24 expression (23). Additionally, a previous study utilizing similar trans-mitochondrial cybrids reported that mitochondria from non-metastatic donors induce a reversal of EMT phenotypes (39). Specifically, the authors used the metastatic osteosarcoma cell line 143B TK^-^ as a nuclear background to mitochondria derived from either epithelial breast cell line MCF10A or TNBC MDA-MB-468. Cybrids with MCF10 mitochondria were significantly less migratory and invasive than either cybrid model with cancerous mitochondria. Tumor growth was also significantly decreased in the 10A/143B cells (39). Therefore, it is possible that OpA targets mitochondria that have undergone the most extensive EMT-associated metabolic reprogramming.

That OpA targets EMT(+) cells through disruption of metabolic pathways is further supported by the dynamic shifts in oxidative and glycolytic metabolism observed in OpA treated cells through metabolomic studies (Sup Fig. 1). Typically, EMT(+) cells utilize glycolysis, even in aerobic conditions, termed the Warburg Effect (75). By upregulating glycolysis, ATP can be produced quickly, and damaging byproducts, such as ROS, are limited (75, 76). Even so, cancer cells still generally have an elevated accumulation of ROS (71, 77). In OpA treated cells, this dependency is challenged, ultimately limiting glycolytic plasticity and decreasing ATP production (Fig. 4). Furthermore, this partial-EMT reversal is also supported by mitochondrial elongation (Fig. 5). Elongation, driven by the promotion of mitochondrial fusion or suppression of fission, has been associated with the oxidative phenotype, while smaller, fragmented mitochondria are observed in more glycolytically active cells (47, 78).

While OpA does increase some parameters of oxidative metabolism, we observed a significant decrease in cellular ROS production in OpA treated cells (Fig. 7). This is likely mediated by decreased GSH transport into mitochondria as OpA binds to the glutathione (GSH) transporter SLC25A40. Therefore, our data suggest that OpA may initially increase ROS levels, inducing oxidative stress and activating antioxidant mechanisms, leading to increased GSH production. Upstream of GSH production, the NRF2 antioxidant response can be activated by oxidative stress, eventually leading to increased expression of several GSH precursors and enzymes via the ARE (57, 59, 61). In support of our hypothesis, we detected NRF2 activation and transcriptional upregulation of genes associated with the ARE enhancer (Fig. 7). As OpA can also reverse certain EMT phenotypes (23), our data suggests OpA limits the ability of cells to compensate for metabolic insults and stress, or metabolic plasticity, a characteristic conferred by EMT.

Based on our findings, we further hypothesize that decreased metabolic plasticity induced by OpA causes compensatory mechanisms, like antioxidation, to be eventually overwhelmed at higher concentrations, leading to metabolic collapse. Indeed, in other studies of OpA in breast cancer, increased cellular ROS abundance was observed in MDA-MB-231 cells at 10 μM, but not 1 μM OpA. Similar to our study, no significant change in ROS abundance was detected in MCF7 cells (18). A different study of OpA in glioblastoma, which also reported elongated mitochondria and involvement of ROS dysregulation, reported direct engagement by OpA with the ROS scavenger NAC and the antioxidant GSH (21). The authors found that OpA interacts with the free thiol groups on these abundant metabolites, forming thia-Michael adducts and decreasing OpA-induced cell death. Interestingly, this result was only obtained when cells were pre-treated, not co-treated, with either NAC or GSH, indicating kinetics may play a significant role in OpA’s mechanism (21). This supports our hypothesis that some metabolic effects can only be captured at higher concentrations (> 10μM) or upon longer incubations of OpA. However, this interaction does not account for the decreased cell death post CDDO pretreatment observed in our data (Fig. 7).

Lastly, OpA has been reported to interact with proteins in complex IV of the ETC in lung cancer (22). Similar to our findings, the authors report decreased membrane potential and ATP production and increased oxidative metabolism. Importantly, this study, utilizing higher concentrations of OpA (1-10 μM) but at lower incubation times, noted a kinetic effect, with ATP production initially spiking before decreasing below the control (22). This supports our hypothesis that OpA may lead to metabolic collapse at higher doses or incubation times. Thus, future studies should focus on first kinetic assays using similar and extended OpA incubation times, then proceed to varying OpA concentrations.

In conclusion, our findings demonstrate that OpA sensitivity is conferred by the mitochondria with metabolic impacts observed at concentrations as low as 50 nM, leading to modulation of metabolic plasticity and EMT characteristics. By engaging with specific targets, OpA challenges EMT conferred metabolic reprogramming, ultimately leading to oxidative stress and activation of compensatory antioxidant mechanisms. Our data suggest OpA-induced cytotoxicity is linked to disrupting EMT-associated metabolic rewiring, causing metabolic stress and eventual collapse at higher concentrations. In the context of breast cancer patients, in particular those with TNBC, the development of a novel agent targeting EMT(+) cells could significantly increase overall survival.

## Declarations

### Ethics Consent Statement

Not applicable

### Data Availability Statement

Data available upon reasonable request. The proteomics raw files have been deposited to PRIDE database. Project accession: PXD062356.

The authors declare no competing financial interests

## Acknowledgements

We thank Sahar Pradhan of the Sayes lab for assistance with the Seahorse instrument, and all the members of the Taube Lab for fruitful discussions.

## Authors’ contributions

Conceptualization: HNP, YT, AK, BC, DR, JHT; Methodology and experimentation: HNP, YT, JT, KLH, SD, EY, AM, NM, JHP, Resources and consulting: SR, BZ, BAK, AB, CMS, AE; Writing—original draft preparation, HNP, JHT; Writing—review and editing, all authors. All authors have read and agreed to the published version of the manuscript.

