## Supplemental Figures for "Ophiobolin A selectively alters mitochondria, metabolism and redox biology in breast cancer and mammary epithelial cells which have undergone epithelial to mesenchymal transition"

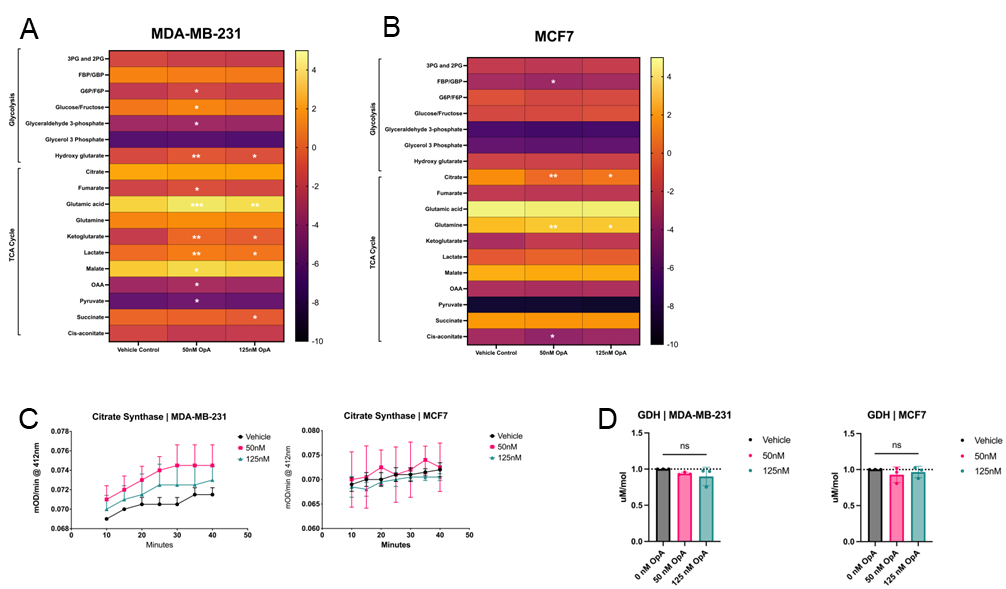

**Supplemental Figure 1**: **OpA Alters Metabolite Abundance in the TCA Cycle and Glycolysis.** Targeted metabolomic analysis via LC-MS in A) MDA-MB-231 and B) MCF7 cells. C) Change in citrate synthase activity over time in MDA-MB-231 (right) and MCF7 (left) cells. D) Activity of glutamate dehydrogenase in MDA-MB-231 (right) and MCF7 (left) cells. All cells treated with either vehicle, 50nM or 125nM OpA for 3h. n = 3. Statistical analysis performed on GraphPad Prism using multiple unpaired t tests and FDR (Q=25%) or 2way-ANOVA with Tukey’s multiple comparison test (*p<0.5; **p<0.01; ***p<0.001; ****p<0.0001).

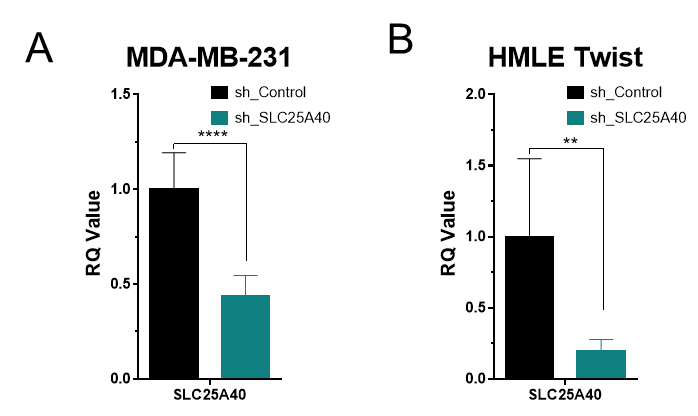

**Supplemental Figure 2 Knockdown of SLC25A40**. mRNA expression of SLC25A40 in A MDA-MB-231or B HMLE-Twist knockdown cells. n = 6. Statistical analysis performed on GraphPad Prism using Student’s t-test (**p<0.01; ****p<0.0001).
